# Novel Evidence for Nitrogen-Dependent Regulation of Nitrate Reductase in Coral Symbiosis

**DOI:** 10.64898/2026.06.15.732369

**Authors:** Chloé Stévenne, Christine Ferrier-Pagès, Renaud Grover, Jean-Christophe Plumier, Stéphane Roberty

## Abstract

The nutritional symbiosis between corals and their photosynthetic dinoflagellate partners underpins the ecological success of reef-building corals in nutrient-poor environments. Although coral holobionts can assimilate the abundant yet highly variable environmental nitrate, direct insight into how nitrate reductase is regulated in these symbiotic algae has been lacking. Here, we provide the first characterization of nitrate reductase protein and gene expression in cultured Symbiodiniaceae exposed to different nitrogen sources and light regimes, revealing the multifactorial nature of its regulation. We demonstrate that nitrate reductase operates as a substrate-induced enzyme: nitrate stimulates protein synthesis in nitrogen-starved cultures, whereas ammonium actively suppresses its expression in a concentration-dependent manner. Light availability and photosynthetic electron transport further modulate protein abundance, suggesting that while nitrate reductase synthesis depends on nitrate availability, its stability may rely on photosynthesis. We also show that nitrate reductase is synthesized within hours of nitrate exposure by symbionts from nitrogen-starved corals, demonstrating that nitrate reduction can occur within the host environment. However, this response is transient and diminished relative to free-living cells, indicating that nitrate reduction is a facultative pathway activated when preferred nitrogen sources are limited. Finally, gene expression measurements and pharmacological inhibition confirm that nitrate reductase regulation is predominantly post-transcriptional, enabling this rapid and reversible control of nitrate assimilation. Together, these findings reveal a tightly regulated and responsive nitrate reduction system in coral symbionts that provides a flexible mechanism contributing to nitrogen homeostasis under fluctuating nutrient regimes.

## 1. Introduction

Coral reefs are among the most biologically diverse and productive ecosystems on Earth, yet they thrive in some of the ocean’s most oligotrophic waters. This paradox is largely resolved by the intimate symbiosis between scleractinian corals and their photosynthetic dinoflagellate of the family Symbiodiniaceae, which, together with other associated microorganisms, constitute a holobiont [1]. Symbiodiniaceae provide their coral hosts with photosynthetically derived carbohydrates, amino acids, and lipids, thereby supporting the host’s metabolism and growth in nutrient-poor environments. In return, the algae receive inorganic nutrients and metabolic waste products from the host and benefit from the protective environment of the symbiosome [2, 3]. While corals are also capable of heterotrophic feeding [4], their survival and ecological success are primarily sustained by their symbionts’ autotrophic productivity [5, 6].

Among the key nutrients required for this partnership, nitrogen is particularly critical due to its role in the synthesis of proteins and other essential biomolecules. In marine environments, nitrogen is present in several forms, with nitrate (NO_3_^-^) being the most abundant source of bioavailable inorganic nitrogen [7, 8]. Coral holobionts preferentially assimilate ammonium (NH_4_^+^) because of its reduced state and the lower energetic cost of assimilation [9, 10]. Both the host and symbionts assimilate NH_4_^+^ via their glutamine synthetase/glutamate dehydrogenase (GS/GDH) and glutamine synthetase-glutamate synthase (GS-GOGAT) pathways. In contrast, coral hosts lack nitrate and nitrite reductase (NR and NiR), preventing them from utilizing NO_3_^-^ directly [11–13].

Nevertheless, due to its environmental abundance, NO_3_^-^ remains a potentially important nitrogen source through assimilation by Symbiodiniaceae [10, 14]. Experimental studies have shown that coral holobionts can remove NO_3_^-^ from the surrounding water [10, 15, 16] and incorporate isotopically labelled nitrate into both algal and host tissues [14, 17, 18]. However, the conditions under which nitrate assimilation occurs, as well as the molecular and enzymatic mechanisms that underpin this process in Symbiodiniaceae, remain largely uncharacterized. Early work using enzymatic assays indicated that NR activity can be modulated by light and nitrate availability, while being suppressed by the presence of ammonium [15, 19]. This led to the hypothesis that host-derived NH_4_^+^ represses NR expression in symbionts [20, 21]. Since then, few studies have directly examined NR in Symbiodiniaceae. Investigations have primarily relied on transcriptomics and gene predictions to examine the expression of nitrate assimilation-related proteins such as nitrate transporters, nitrate and nitrite reductase or GS-GOGAT proteins [22–25]. As such, little is known about how environmental conditions—including nitrogen source, light, and circadian rhythms—influence NR expression and activity at transcriptional or post-translational levels.

Furthermore, the intracellular location of symbionts poses an additional layer of complexity for nitrate acquisition. At physiological pH, NO₃⁻ is charged and thus requires active transport across lipid membranes [21]. While nitrate transporters (NRT) have been identified in Symbiodiniaceae [23, 26–28], the mechanism by which nitrate is transported across the host cell membrane into the symbiosome remains unclear. The only indication of a potential transport pathway is the identification of SIALIN-like sequences in the genomes of several coral species, possibly homologous to the H^+^:2NO_3_^-^ cotransporter SIALIN [29].

Taken together, these knowledge gaps highlight the need for a deeper understanding of nitrate assimilation pathways in coral symbioses, both at the molecular and cellular levels. Elucidating these mechanisms is essential for understanding how corals manage nitrogen resources, particularly under changing environmental conditions that may alter nutrient availability. In this study, we investigated the expression patterns of nitrate reductase in Symbiodiniaceae, both in culture and *in hospite*. Using western blotting (WB) and qRT-PCR, we characterized NR protein and gene expression in response to different nitrogen sources (NO_3_^-^ and NH_4_^+^) and examined the influence of light and photosynthetic electron flow. In addition, we tested the effects of various inhibitors to shed light on potential regulatory mechanisms at transcriptional, post-transcriptional and post-translation levels. Finally, we examined NR protein expression in symbionts from three coral species following NO_3_^-^enrichment, providing new insight into nitrate assimilation within the coral-algal symbiosis.

## 2. Materials and Methods

### 2.1. Symbiodiniaceae Strains and Culture Conditions

Symbiodiniaceae strains belonging to *Breviolum minutum* (SSB01) and *Symbiodinium microadriaticum* species were used for culture-based experiments. SSB01 was originally isolated from the sea anemone *Exaiptasia diaphana* line H2 [30], while *S. microadriaticum* was isolated from a colony of the Red Sea coral *Stylophora pistillata*. Both strain identities were confirmed by ITS2 sequencing as described in ***Supporting Information S1***.

Symbiodiniaceae cultures were grown and maintained in a Percival I-36LLX incubator (Percival Scientific Inc, USA) at 26°C and under 150 µmol photon m^-2^ s^-1^ on a 12h:12h light:dark cycle provided by cool white 10,000 K (Sylvania Aquastar F18W/174-T8) and blue 20,000°k lights (Aquavie Lumivie SM T8). Symbiodiniaceae cells were maintained in a modified f/2 medium (***Supporting Information S2-S4***) [31, 32] made in artificial seawater (ASW) at a salinity of 34 prepared with Coral Pro Salt (Red Sea Aquatics, UK). Each strain was grown with 500 µM of nitrogen either in the form of nitrate (NO_3_^-^) or ammonium (NH_4_^+^) and were kept in triplicate culture flasks for each condition in volumes ranging from 150 mL to 300 mL. Exponential growth was maintained by subculturing into fresh medium once a week. All experiments were conducted during the exponential growth phase of the cultures.

### 2.2. Experimental treatments applied to Symbiodiniaceae in cultures

The five experimental treatments conducted to elucidate the factors influencing NR expression in Symbiodiniaceae are detailed in ***Fig. 1***. Before treatments, cultures grown under long-term (LT) conditions in 500 µM NO_3_^-^ f/2 medium were transitioned to a nitrogen-free (NF) f/2 medium for 4 days to deplete cellular nitrogen levels. This approach was adopted following observations that NR protein was not detectable in nitrogen-replete algae (***Supporting Information S5***).

**Fig. 1.**
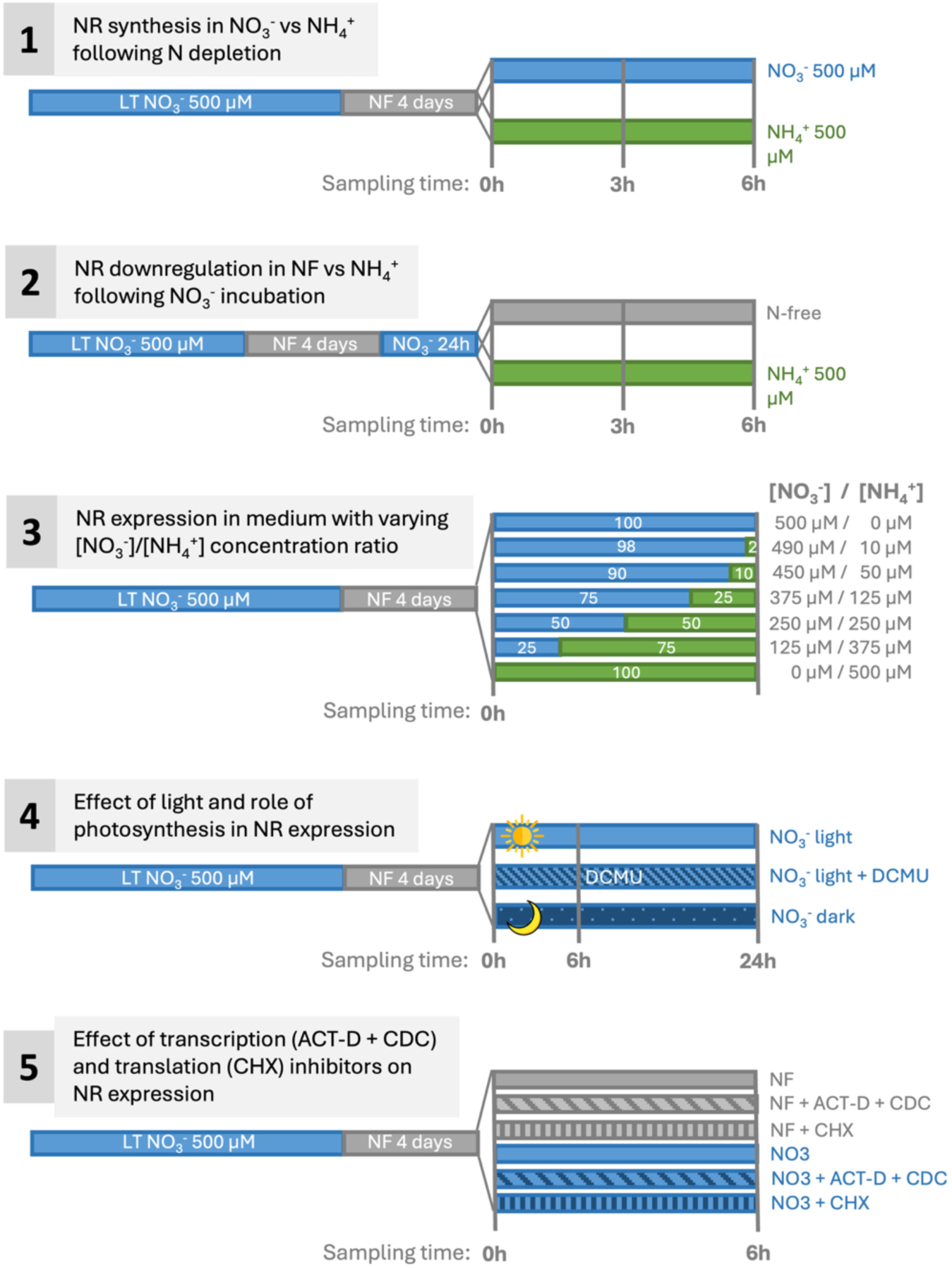
Summary of experimental conditions of the 5 experiments investigating nitrate reductase expression and regulation in Symbiodiniaceae in cultures. Treatment conditions are schematically represented by colour bands corresponding to the incubation interval and the nitrogen treatment. The colours are associated with the nitrogen treatment with grey for nitrogen-free, blue for nitrate and green for ammonium. Special treatments such as the addition of inhibitors and light treatments are represented by patterns on the bands. The total nitrogen concentration in treatment media was always set to 500 µM. Experiments 1,2 and 3: Symbiodiniaceae cells were transitioned between the different nitrogen treatments (NF, NO_3_^-^, NH_4_^+^ and mixtures of NO_3_^-^ and NH_4_^+^) and incubated for 3 to 6 h. Experiment 4: 12h dark-incubated (night cycle) Symbiodiniaceae were transitioned under one of three conditions and incubated for 6 to 24 h: control light (150 µmol photons m^-2^ s^-1^), complete darkness (0 µmol photons m^-2^ s^-1^), or standard light with 40 µM of 3-(3,4-dichlorophenyl)-1,1-dimethylurea (DCMU), a photosystem II (PSII) inhibitor. Experiment 5: Symbiodiniaceae cells were placed under six experimental conditions and incubated for 6 h. These conditions combined two nitrogen treatments–NF or NO_3_^-^ 500 µM–with three inhibitor treatments: no inhibitors, transcription inhibitors, or translation inhibitors. Transcription inhibitors actinomycin-D (ACT-D) and cordycepin (CDC) were added together at final concentrations of 0.5 µM and 20 µg/mL respectively [33], while translation inhibitor cycloheximide was added at a final concentration of 10 µg/mL [34, 35].

Experiments 1 and 2 aimed to identify the individual effects of the nitrogen source on NR expression, while Experiment 3 examined NR expression in response to different concentration ratios of NO_3_^-^ and NH_4_^+^. Experiment 4 investigated the influence of light and photosynthesis on NR protein and gene expression. Experiment 5 explored the role of transcriptional and post-transcriptional regulation on NR expression using the transcription inhibitors actinomycin-D (ACT-D) and cordycepin (CDC), and the translation inhibitor cycloheximide.

### 2.3. NR protein expression in cultured Symbiodiniaceae

At each sampling time (see ***Fig. 1***), 15 mL of Symbiodiniaceae culture (15-30 mg fresh weight) were collected per replicate. Protein extracts were prepared using a Tris-base extraction buffer with a protease inhibitor cocktail and cell disruption using 250 µL of glass beads (710-1180 nm, acid-washed, Sigma-Aldrich) in a TissueLyser II (QIAGEN) for 10 min at 30 Hz. Protein concentrations were determined by spectrophotometry using the colorimetric RC DC^TM^ Protein Assay (BioRad, USA).

The protein extracts were then prepared for western blotting together with a sample of purified nitrate reductase isolated from *Arabidopsis thaliana* (N0163-1VL, Sigma-Aldrich, Merck, Germany), which was used as a positive control. Denatured protein extracts in Laemmli buffer [36] were separated by electrophoresis on an SDS-PAGE gel (80V for 30 min followed by 140V for 60 min) and transferred to a polyvinylidene difluoride membrane (Immun-Blot® PVDF, Bio-Rad, USA). Primary antibodies for nitrate reductase detection (dilution 1/1000, Assimilatory NR, Agrisera AS08 310) were incubated first and primary antibodies for Cyt f detection (dilution 1/5000, algal cytochrome f PetA of the thylakoid Cyt b6/f-complex, Agrisera AS06 119), used as a protein loading control, were incubated the next day after a first visualization of NR proteins.

Immunoreactive bands were detected with horseradish peroxidase (HRP) conjugated to anti-rabbit secondary antibodies (dilution 1/25000, goat anti-rabbit IgG HRP conjugated, Agrisera AS09 602) and using the Amersham ECL Prime detection system (Cytiva, USA) on an iBright^TM^ FL1000 imaging system (Thermo Scientific, USA). Analysis and quantification of immunoblot bands were performed using the iBright^TM^ analysis software (Thermo Scientific, USA). Nitrate reductase protein expression was calculated as the ratio between NR band and Cyt f band signal expressed as local background corrected volume (full method in ***Supporting Information S6***).

### 2.4. NR gene expression in cultured Symbiodiniaceae

Nitrate reductase gene expression was investigated in SSB01 samples from experiments 1, 2, 4 and 5 (***Fig. 1***) using quantitative RT-PCR. At each sampling time, 15 mL of SSB01 culture (15-30 mg fresh weight) was sampled from each replicate in each experimental condition. RNA was extracted from the algae using the Purelink® RNA Mini Kit (Thermo Scientific, USA) according to the manufacturer’s instructions. Cell disruption was achieved in 0.9 mL of lysis buffer with acid-washed glass beads (710-1180 µm, Sigma-Aldrich) using bead-beating on the TissueLyser II (QIAGEN) for 7 min at 30 Hz. RNA was precipitated with absolute molecular grade ethanol, purified on spin columns including DNase treatment and eluted in 25 µL RNase-free water. Total RNA concentrations were quantified, and RNA quality was assessed by spectrophotometry on a Take3 microspot plate using a Synergy Mx BioTek^TM^ plate reader. cDNA was synthetized from 0.25-1 µg RNA (varying by experiment but consistent within each) by reverse transcription using the RevertAid H Minus First Strand Kit (Thermo Scientific, USA), with a 60 min incubation at 42°C followed by 5 min at 70°C.

The primer pairs Cyc_Maru [37] and Act1_May [38] for genes cyclophilin and actin1, respectively, were selected as housekeeping genes (HKG) (***Supporting Information S7***). The nitrate reductase primer pair NR_s6_34 was selected from Xiang et al. (2020) based on the putative NR transcript s6_34 [23]. This primer pair was successfully used by Maruyama et al. (2022) to study NR gene expression levels in SSB01 cultures and in symbionts isolated from the sea anemone *Exaiptasia diaphana* [37]. Primer sequences and specifications are summarized in ***Table 1***.

**Table 1.**
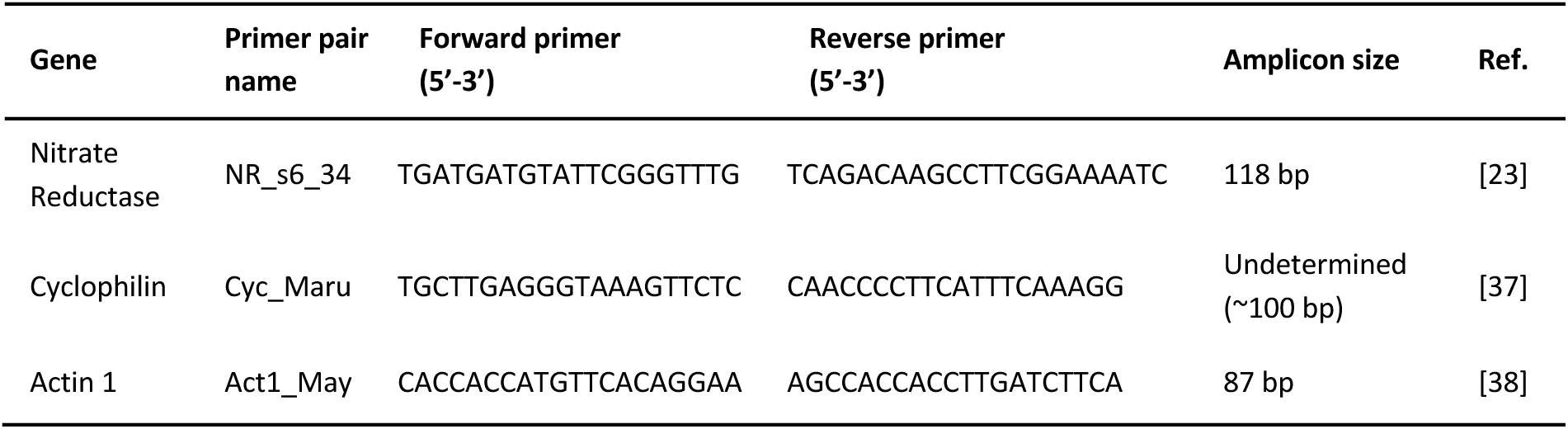
Primer pair names, sequences, and amplicon sizes for gene for quantitative real-time PCR of gene interest (NR) and housekeeping genes (Cyc and Act1) in SSB01 (*Breviolum minutum*). Primer sequences and amplicon sizes were obtained from the literature.

Real-time polymerase chain reactions (qPCRs) were performed in technical triplicate for each sample using 4 µL of 1:50 diluted templates in 10 µL reaction volumes with Takyon^TM^ Low ROX SYBR 2X MasterMix blue dTTP (Eurogentec, Belgium). qPCRs were performed on a QuantStudio^TM^ 5 system (Thermo Scientific, USA) according to a PCR profile consisting of an initial incubation of 2 min at 50°C and 2 min at 95°C, followed by 40 cycles of 15 s at 95°C, and 1 min at 60°C. Runs were terminated with melt curves from 60°C to 95°C and RNase-free water controls were included as negative controls. Relative expression of genes was expressed as ΔΔCt and fold gene expression as 2^-ΔΔCt^ (***Supporting Information S8***). Statistical analyses were performed on ΔΔCt values using ANOVA and *post-hoc* Tuckey’s honestly significant difference (HSD) tests.

### 2.5. NR expression in *in hospite* Symbiodiniaceae

The nitrate reductase expression by Symbiodiniaceae *in hospite* was investigated in *Stylophora pistillata* (Esper 1792), *Turbinaria reniformis* (Bernard 1896) and *Echinopora lamellosa* (Esper 1791), and their freshly isolated symbionts. These three coral species from different families were selected for their association with diverse symbiont assemblages [39–44] and their different trophic and nutritional capabilities [44–47]. Coral colonies were maintained in flow-through seawater aquaria at the Centre Scientifique de Monaco (Monaco). Nubbins were produced from mother colonies, evenly distributed between two independent tanks, and allowed to recover under controlled environmental conditions. During the four-week recovery, they were fed twice weekly with Artemia nauplii, and feeding was stopped two weeks before experiments to prevent interactions with nitrogen enrichments (details in ***Supporting Information S9***).

Two experiments were performed to investigate NR protein expression in *in hospite* Symbiodiniaceae (experiment A performed on all 3 coral species) and in Symbiodiniaceae freshly isolated (FIS) from their coral host (experiment B performed on FIS from *A. pistillata* and *T. reniformis*) (***Fig. 2***). For both experiments, coral nubbins were first incubated in nitrogen-depleted water for 24 h (undetectable levels of dissolved inorganic nitrogen, DIN).

**Fig. 2.**
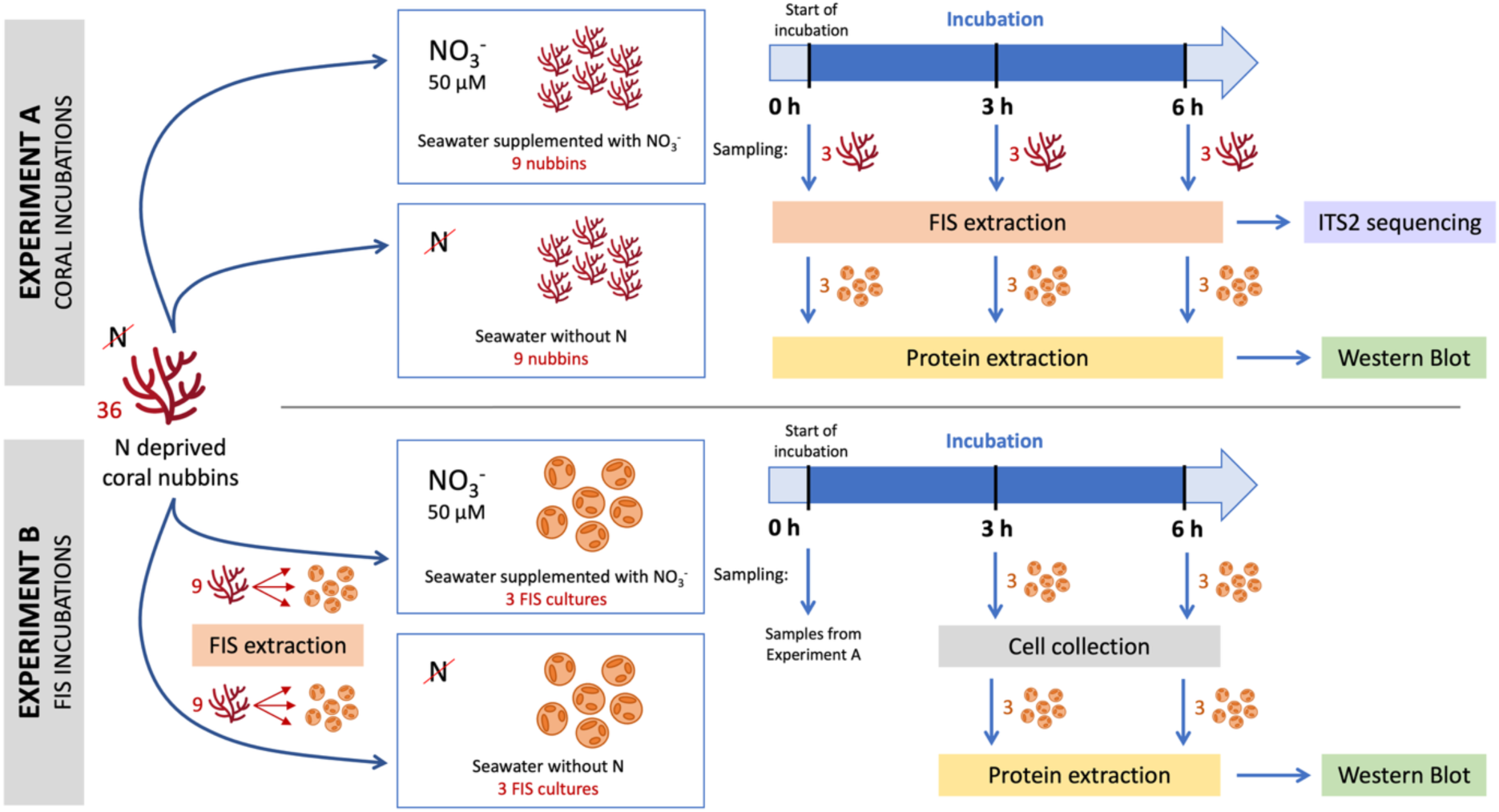
Schematic representation of experimental conditions of the 2 experiments investigating nitrate reductase protein expression in *in hospite* symbionts of the corals *S. pistillata*, *T. reniformis*, and *E. lamellosa* (***Experiment A***), and in freshly isolated symbionts (FIS) from the corals *S. pistillata*, *T. reniformis* (***Experiment B***). **A.** All treatments were carried out in duplicate 28 L tanks with water temperature maintained at 25°C ± 0.5 °C, with constant agitation provided by water pumps, and under 150 µmol photons m^-2^ s^-1^ in a 12h:12h day:night cycle. All experimental tanks were kept as a closed system and nitrate enrichment was achieved by mixing a NaNO_3_ concentrated stock solution to the tank seawater just before coral nubbins were transferred to the experimental tanks. **B.** FIS were incubated in 200 mL culture flasks with ventilated cap, at a temperature of 25°C ± 0.5 °C with gentle agitation using a magnetic stirrer and under 150 µmol photons m^-2^ s^-1^ constant light.

For experiment A, 6 nubbins per coral species were transferred from N-depleted seawater into each treatment. Control nubbins were incubated for 6 h in natural oligotrophic filtered seawater (FSW), while treated nubbins were incubated in 50 µM nitrate-enriched FSW. Coral nubbins were sampled in triplicate for each coral species at 0 h (prior to incubation in the experimental treatments), at 3 h and 6 h after being incubated in treated water. Samples were immediately placed in liquid nitrogen for preservation at -80°C until downstream processing for western blotting.

For experiment B, 18 N-depleted nubbins per coral species were sampled. Symbionts were freshly isolated (FIS) from the coral host tissues (details in ***Supplementary Information S9***). Isolated symbionts from 3 nubbins of each species were pooled to constitute biological replicates (3x) of each treatment. Control FIS were incubated for 6h in 70 mL of natural oligotrophic FSW, while treated FIS were incubated in 70 mL of natural FSW supplemented with 50 µM NO_3_^-^. After 3 and 6 h of incubation, 10 mL of FIS culture were sampled and immediately processed for protein extraction.

All samples were processed for protein extraction, quantification, and western blotting as described above for cultured Symbiodiniaceae. FIS samples were preserved in RNAlater® solution (Thermo Scientific, USA) until downstream analysis for ITS2 sequencing as described in ***Supporting Information S1***.

### 2.6. Statistical analyses

Statistical analyses were performed in R (R version 4.3.1) [48]. Data were checked for normality and homoscedasticity. If normality and/or homoscedasticity conditions were not met, data were log-transformed (using base 10 logarithms). A 2-way (time x treatment) repeated measure ANOVA (rmANOVA) was applied to analyse the responses in experiments 1 and 2, as well as the experiments investigating NR expression in *in hospite* symbionts (experiments A and B). To achieve this, linear mixed-effect (LME) models (lme() function, nlme R package, v3.1.163) were fitted to the data to investigate the response of NR protein and gene expression (dependent variable) to treatment (NO_3_^-^ and NH_4_^+^) and time (fixed effects, independent and interacting), while accounting for repeated measures on replicate algal cultures (random effect). Similarly, LME was used for experiment 5 to investigate the effects of treatment (light, dark and light + DCMU) and time on both NR protein and gene expression. One-way ANOVA was applied to the western blot data from experiment 4 to test the significance of treatment (varying [NO_3_^-^]/[NH_4_^+^] ratio) on NR protein expression. For experiments 3 and 6, two-way ANOVAs were used to analyse the independent and interacting effects of nitrogen treatment and inhibitor treatments on NR protein expression. When significant effects were observed, post-hoc Tukey HSD tests were performed. Differences were considered significant when p < 0.05.

## 3. Results

### 3.1. Effect of nitrogen source on NR expression

Our initial immunodetection assays revealed little to no detectable nitrate reductase in protein extracts from Symbiodiniaceae cultured under nitrogen-replete conditions or isolated from nitrogen-replete corals (***Supporting Information S5***). Therefore, all subsequent experiments in this study included a nitrogen depletion step prior to the various treatments. We first identified the individual effects of the nitrogen source on NR expression. Nitrogen-depleted *B. minutum* were incubated with 500 µM NO₃⁻ or NH₄⁺, and NR protein and gene expression were monitored at 0, 3, and 6 hours. Both nitrogen source and time significantly influenced NR protein levels (***Fig. 3A***; 2-rmANOVA, p < 0.0001). NR protein, initially undetectable, appeared at 3 h and increased further at 6 h with NO₃⁻ (Tukey HSD: p < 0.001). In contrast, NR remained undetectable under NH₄⁺. These results were confirmed in *S. microadriaticum* (***Supporting Information S10***). We next assessed NR protein degradation. NR-expressing cells were incubated in NF or NH₄⁺ media. Both time and treatment significantly affected NR expression (***Fig. 3B***; 2-rmANOVA, p < 0.01). After 3 h, NR was stable in NF but decreased significantly with NH₄⁺ and after 6 h, NR was undetectable in both conditions (Tukey HSD, p < 0.001). In both experiments, NR gene expressions remained unchanged regardless of nitrogen treatment or incubation time (***Fig. 3C,D***).

**Fig. 3.**
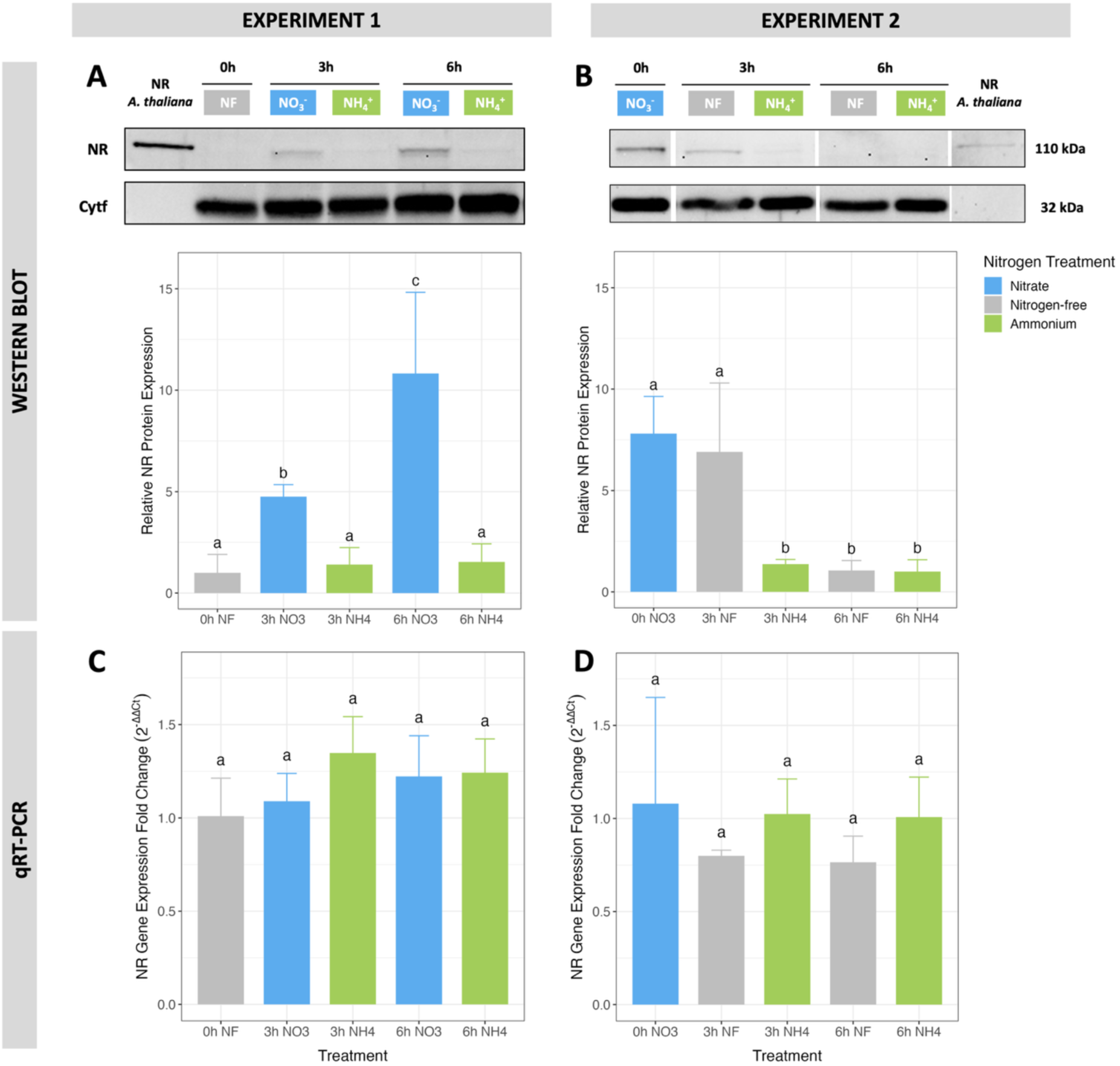
Kinetics of NR expression in *B. minutum* SSB01. Experiment 1: NR protein (**A**) and gene (**C**) expression after incubation of N-depleted SSB01 in f/2 medium supplemented with 500 µM NO_3_^-^ or 500µM NH_4_^+^ for 3 and 6 h. Experiment 2: NR protein (**B**) and gene (**D**) expression following transfer of NR-expressing cells (pre-incubated 24 h in 500 µM NO_3_^-^) to nitrogen-free (NF) or 500 µM NH_4_^+^ for 3 and 6 h. Each western blot (WB) shows a representative membrane (out of 3 biological replicates), with NR protein (∼110 kDa) and the loading control Cytf (∼32 kDa). Mean relative WB signals intensities from all replicates are shown in the bar plots below. qRT-PCR results are represented as fold changes in NR gene expression (2^-ΔΔCt^) relative to time 0. Error bars represent 95% confidence intervals (CI) and statistically significant differences are indicated by different letters (Tukey HSD, p < 0.05).

In experiment 3, we explored how varying NO₃⁻/NH₄⁺ ratios affected NR expression. Algae were incubated for 6 h with 500 µM total nitrogen using NO₃⁻/NH₄⁺ ratios of 100–0, 98–2, 90–10, 75–25, 50–50, 25–75, and 0–100%. NR protein levels declined significantly with decreasing NO₃⁻/NH₄⁺ ratios (ANOVA, p < 0.0001), though the effect was non-linear (***Fig. 4***). For example, 25% NH₄⁺ reduced NR expression to 60 ± 5% of the 100% NO₃⁻ value, while 50% NH₄⁺ led to 28 ± 9%, similar to levels in NF (16 ± 7%) and 100% NH₄⁺ (24 ± 12%) conditions (Tukey HSD, p > 0.05).

**Fig. 4.**
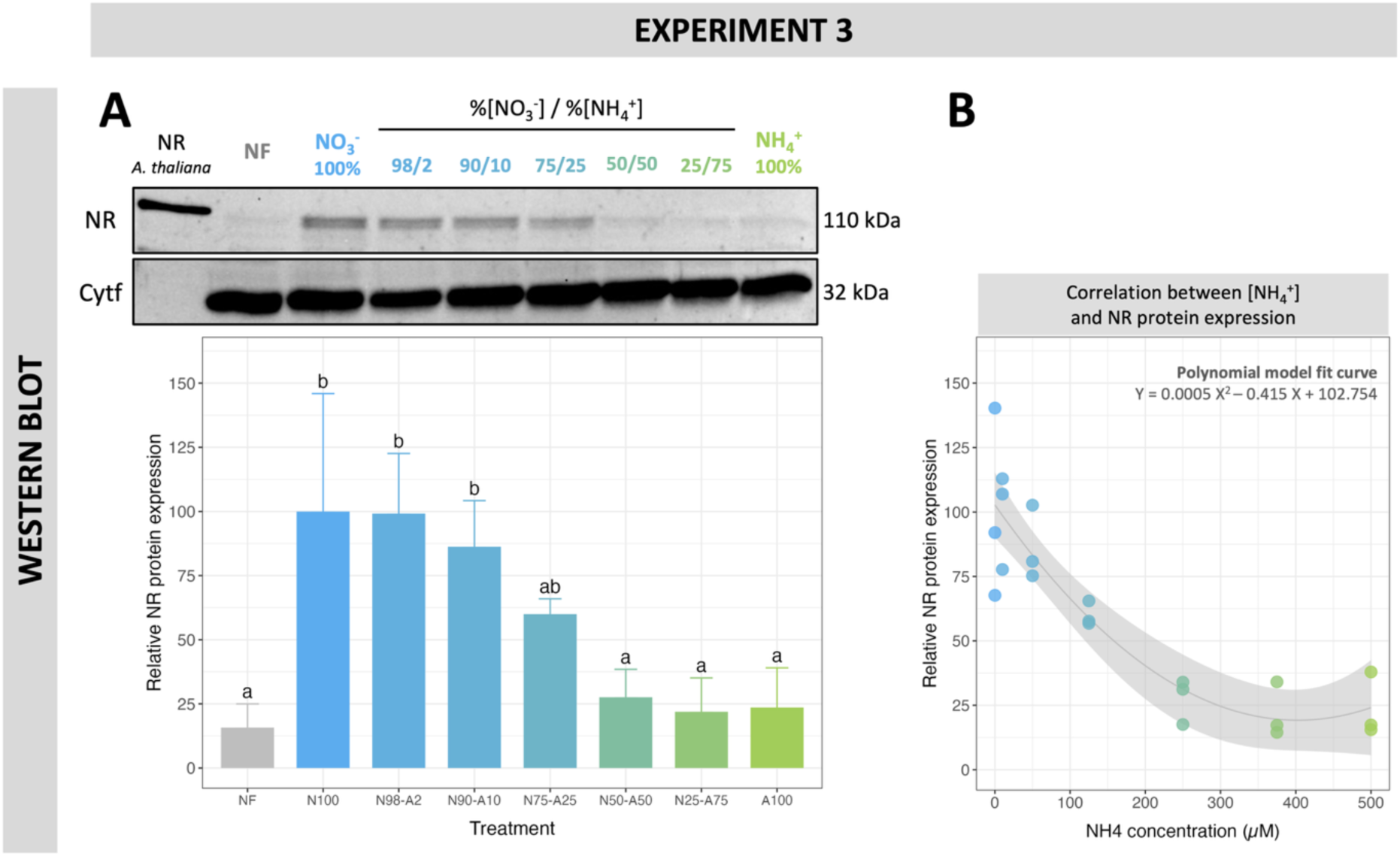
Effect of [NO_3_^-^]/[NH_4_^+^] concentration ratios on NR expression in *B. minutum* SSB01. N-depleted SSB01 were incubated for 6 h in F/2 media containing 500 µM total nitrogen with varying NO_3_^-^/NH_4_^+^ concentrations ratios. **A.** Representative WB membrane (out of 3 replicates) is shown. The upper band (∼110 kDa) corresponds to the NR protein; the lower band (∼32 kDa) to the loading control (Cytf). Mean relative WB signal intensities (n=3) are shown in the bar plot below. Statistically significant differences are indicated by different letters. Error bars correspond to the 95% CI. **B.** Scatter plot of relative NR expression as a function of NH_4_^+^ concentration, with a fitted polynomial regression (grey line) and 95% CI (grey band).

### 3.2. Effect of light on NR expression

The next experiment aimed to investigate the contribution of light on NR expression. Nitrogen-depleted SSB01, dark-acclimated for 12 h, were incubated with 500 µM NO₃⁻ for 6 and 24 h under three light conditions: light (150 µmol photons m⁻² s⁻¹), light with 40 µM DCMU (an inhibitor of PSII electron flow; [49, 50]), and complete darkness. Both light treatment and incubation time significantly affected NR protein expression (***Fig. 5A***; 2-way ANOVA, p < 0.0001). As in previous experiments, NR protein expression increased significantly after 6 h in light (Tukey HSD, p < 0.0001). The protein was also synthetised in darkness (p < 0.05), although at lower levels than in light (p < 0.0001) or light + DCMU (p = 0.01). After 24 h, NR expression in darkness had decreased to levels comparable to nitrogen-depleted cells and remained significantly lower under light + DCMU than under light (p = 0.0001).

**Fig. 5.**
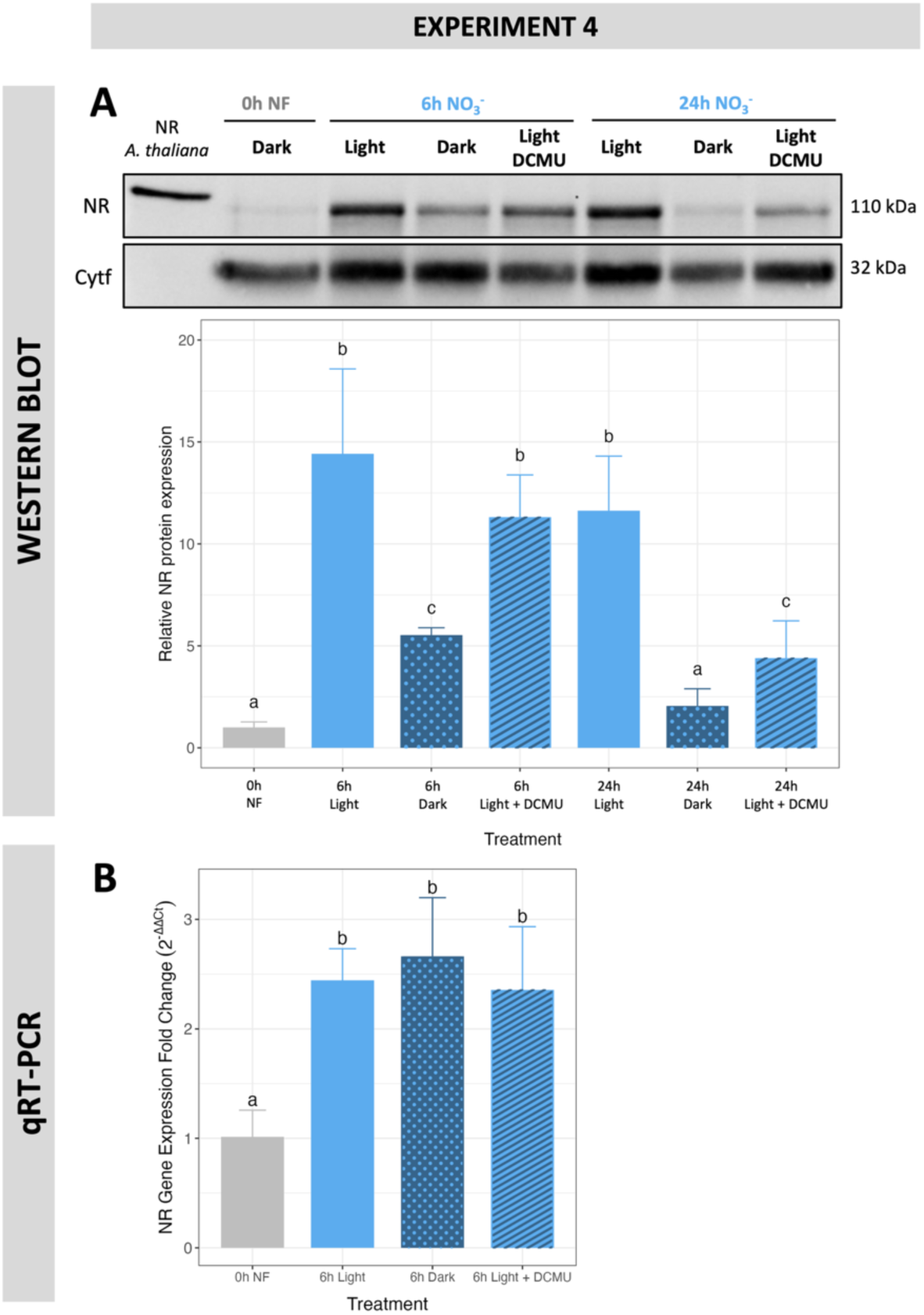
Role of light and photosynthesis in NR expression in *B. minutum* SSB01. N-depleted SSB01 were incubated for 6 and 24 h in f/2 medium with 500 µM NO_3_^-^ under three light conditions (light, dark, and light + 40 µM DCMU). **A.** Representative WB membrane (out of 3 replicates), the upper band (∼110 kDa) corresponds to the NR protein; the lower band (∼32 kDa) to the loading control (Cytf). Mean relative band intensities from all replicates (n=3) are shown in the bar plot. **B.** qRT-PCR results show NR gene expression fold change (2^-ΔΔCt^) relative to time 0. Error bars indicate 95% CI. Different letters indicate significant differences (Tukey HSD,p < 0.05).

NR gene expression, measured after 6 h of NO₃⁻ incubation, was significantly higher in NO₃⁻-treated cells than in nitrogen-depleted controls (ANOVA, p < 0.0001), but was not affected by the light treatments (***Fig. 5B***).

### 3.3. Effect of transcription and translation inhibitors on NR expression

Given the lack of correlation between NR protein and gene expression observed in Experiments 1, 2, and 4, Experiment 5 examined the roles of transcriptional and post-transcriptional regulation using specific inhibitors of transcription (actinomycin D, ACT-D; and cordycepin, CDC) and translation (cycloheximide, CHX). ACT-D binds DNA and CDC disrupts mRNA polyadenylation [51, 52], while CHX blocks protein elongation [53]. Both nitrogen-free (NF) and NO₃⁻ conditions were used, with the former serving as a control due to naturally lower NR protein levels.

Nitrogen level and inhibitor treatment each had a significant effect on NR protein expression (2-way ANOVA: p < 0.01), with no interaction effect, indicating consistent responses across conditions (***Fig. 6A***). NR protein remained low in NF and NF + ACT-D/CDC, similar to the nitrogen-depleted and NF + CHX treatments (Tukey HSD, p > 0.05). In contrast, NR protein expression increased significantly in NO₃⁻ and NO₃⁻ + ACT-D/CDC (p > 0.01), but was suppressed in NO₃⁻ + CHX, to levels indistinguishable from NF (p > 0.05). *NR* gene expression was not affected by nitrogen or inhibitor treatments (2-way ANOVA, p > 0.05 for all effects) (***Fig. 6B***).

**Fig. 6.**
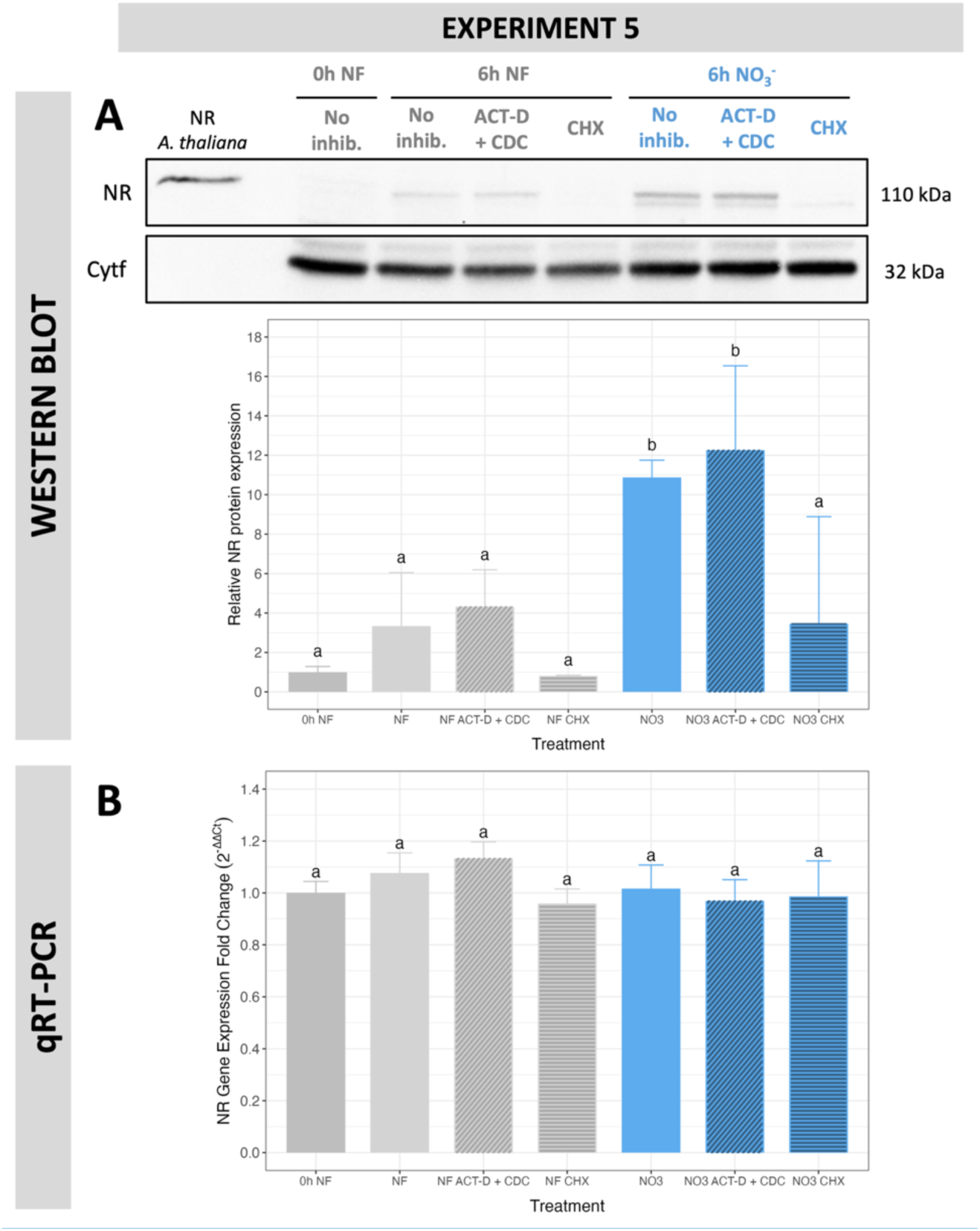
Effect of transcription and translation inhibitors on NR expression in *B. minutum* SSB01. N-depleted SSB01 were incubated in nitrogen-free f/2 NF or 500 µM NO_3_^-^, with or without transcription inhibitors (ACT-D + CDC), or translation inhibitor cycloheximide (CHX). **A.** Representative WB membrane (out of 3 biological replicates), the upper band (∼110 kDa) corresponds to the NR protein; the lower band (∼32 kDa) to the loading control (Cytf). Bar plot shows mean relative WB signal from all replicates (n = 3). **B.** qRT-PCR results expressed as NR gene fold change (2^-ΔΔCt^) relative to time 0. Error bars indicate 95% CI. Different letters denote significant differences (Tukey HSD, p < 0.05).

### 3.4. Expression of NR in coral symbionts

Building on previous experiments in *B. minutum* and *S. microadriaticum* under NO₃⁻-enriched culture conditions, we next examined NR protein expression in symbiotic contexts. Experiments A and B assessed NR expression in *in hospite* (coral tissue) and freshly isolated symbionts (FIS), respectively, using three coral species with known ITS2 symbiont profiles. *S. pistillata* predominantly hosted type A1 Symbiodiniaceae (*S. microadriaticum*, 98%), while *T. reniformis* and *E. lamellosa* mainly harbored *Cladocopium* (86% and 64%), with *E. lamellosa* also containing *Durusdinium* (25%).

Preliminary tests showed no NR protein in *S. pistillata* symbionts under nitrate-enriched conditions without prior nitrogen depletion (***Supporting Information S11***). Thus, in subsequent incubations, corals were first nitrogen-starved, then exposed to either natural filtered seawater (FSW) or FSW enriched with 50 µM NO₃⁻, following protocols shown to induce NR expression in cultured Symbiodiniaceae.

NR protein was detected *in hospite* via western blot in all three species after 3 h incubation in both control and NO₃⁻-enriched seawater (***Fig. 7A,C,E***). Expression tended to be higher under NO₃⁻ but only reached statistical significance in *E. lamellosa* (Tukey HSD, p = 0.016), due to high variability and low detection levels in *S. pistillata* and *T. reniformis*. In all cases, NR levels were significantly lower after 6 h compared to the 3 h NO₃⁻ condition (p < 0.05).

**Fig. 7.**
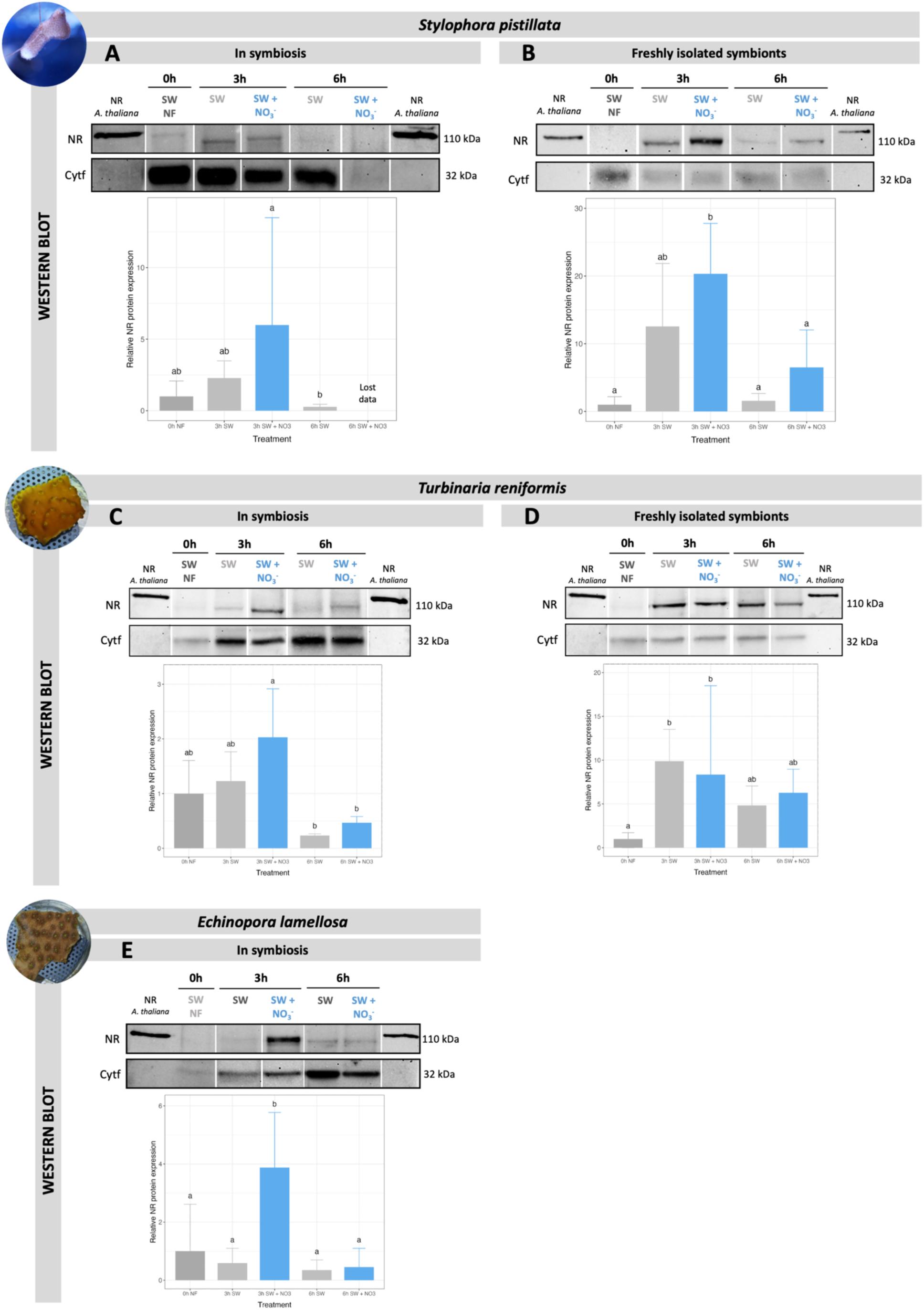
NR protein expression in symbionts from three coral species, in hospite and after isolation. Western blot analysis of NR protein expression in symbionts from N-depleted corals, *S. pistillata* (**A**), *T. reniformis* (**C**) and *E. lamellosa* (**E**), incubated for 3 and 6 h in either seawater (SW) or SW enriched with 50 µM NO_3_^-^. NR expression in freshly isolated symbionts (FIS) from *S. pistillata* (**B**) and *T. reniformis* (**D**) under the same incubation conditions. For clarity, only one representative WB band per condition (from a total of 3 biological replicates) is shown. The upper band (∼110 kDa) corresponds to the NR protein; the lower band (∼32 kDa) to the loading control (Cytf). Bar plots below each blot show mean relative NR signal intensities (n = 3). Error bars represent the 95% confidence interval (CI), and different letters indicate statistically significant differences (Tukey HSD, p < 0.05).

In FIS, NR protein was detected in both FSW and NO₃⁻ conditions after 3 and 6 h in symbionts from *S. pistillata* and *T. reniformis* (***Fig. 7B,D***). While differences were not statistically significant due to high variability (p > 0.5), NR expression in *S. pistillata* FIS was consistently higher under NO₃⁻, a pattern not observed in *T. reniformis*.

## 4. Discussion

Nitrate is the most abundant form of DIN in coral reef environments but exhibits strong temporal and spatial variability due to both natural processes and anthropogenic inputs [7, 54, 55]. Understanding how nitrate is assimilated by the coral holobiont is therefore crucial for elucidating nitrogen cycling and coral resilience. Since Crossland and Barnes (1977) first reported nitrate reductase activity in Symbiodiniaceae [15], little attention has been given to NR protein expression itself. Most subsequent studies have inferred nitrate assimilation from uptake measurements at the holobiont level, generally assuming that symbiont NR activity underpins this process [9, 10, 16, 56]. Here, we provide the first direct evidence of NR protein and gene expression in Symbiodiniaceae in response to different nitrogen sources, revealing how this enzyme is regulated at multiple levels.

### Regulation of NR by nitrogen source and NH₄⁺ inhibition

Our results demonstrate that NR behaves as a substrate-induced enzyme: nitrate induces its synthesis in nitrogen-starved Symbiodiniaceae, whereas ammonium does not (***Fig. 3A,B***). This pattern, consistent across both *B. minutum* and *S. microadriaticum*, parallels findings in higher plants, chlorophytes, and diatoms, where NR expression is tightly regulated by nitrate availability and repressed by more reduced nitrogen forms [57]. We further observed that in the presence of NH₄⁺, NR protein disappears more rapidly (within 3 h) than under nitrogen starvation alone (***Fig. 3B***), indicating that ammonium exerts a direct inhibitory effect on enzyme expression. Previous studies have reported lower NO_3_^-^ uptake by Symbiodiniaceae [9, 22, 58] and coral holobionts [10, 59], as well as the downregulation of most nitrate transporter (NRT) genes in the presence of NH_4_^+^ [22]. By combining these previous observations with our results, we deduce that ammonium acts as a potent suppressor of nitrate assimilation at both transport and enzymatic levels.

NR synthesis in free-living Symbiodiniaceae is further modulated by the relative availability of nitrate and ammonium. When supplied together, NR expression declined sharply beyond a threshold ratio of NH_4_^+^ to NO_3_^-^ (***Fig. 4***), suggesting a switch-like response. Equimolar concentrations were sufficient to fully repress NR, consistent with earlier findings that NO_3_^-^uptake ceases when NH_4_^+^ constitutes >25% of total nitrogen [58]. These results indicate that ammonium exerts a concentration-dependent, active repression of NR expression, reflecting a hierarchy in nitrogen source preference.

### Post-transcriptional and post-translational regulation

Whether ammonium represses NR synthesis or promotes its degradation remains unclear. In chlorophyte algae, both processes have been implicated [60], but regulatory pathways in dinoflagellates are still poorly understood. Given their unusual genomic organization–tandem gene arrays, permanently condensed chromosomes, and a paucity of canonical transcription factors [61–64]–Symbiodiniaceae are thought to rely heavily on post-transcriptional control. Consistent with this, NR mRNA levels remained stable across treatments (***Fig. 3***), indicating that NR regulation occurs primarily at or after translation. Similar decoupling of transcript and protein abundance has been reported in the Symbiodiniaceae *Fugacium kawagutii* [22].

Pharmacological inhibition experiments support this interpretation: NR protein levels declined with cycloheximide (CHX), which blocks translation [53], but remained unaffected by the transcriptional inhibitors actinomycin-D (ACT-D) and cordycepin (CDC) [51, 52] (***Fig. 6***). Together, these findings indicate that NR expression in Symbiodiniaceae is primarily regulated at the translational level. This is consistent with emerging evidence of pervasive post-transcriptional mechanisms in dinoflagellates, including small RNA-mediated regulation and RNA editing via pentatricopeptide repeat (PPR) proteins [65–69]. The constitutive stability of NR transcripts likely enables rapid enzyme synthesis when nitrate becomes available, which could constitute an adaptive trait for organisms experiencing fluctuating nitrogen regimes.

Post-translational regulation may provide an additional control layer. In higher plants, NR activity is modulated via phosphorylation-dependent binding of 14-3-3 proteins, leading to reversible inactivation [70], or to irreversible inhibition by protein binding, and targeted proteolysis [71]. Although such mechanisms have not yet been described in dinoflagellates, similar processes could fine-tune NR activity in response to environmental cues.

### Light-dependent regulation

Nitrate assimilation is energetically linked to photosynthesis, as NR requires photosynthetically derived electrons to reduce nitrate [72]. While light-dependent NR activity is well established in plants and algae [57, 73–75], evidence in Symbiodiniaceae and coral holobionts has been inconsistent [9, 10, 15, 16, 76]. Our results show that protein levels decline in darkness and are partially suppressed by DCMU, implicating PSII-derived electron flow in maintaining NR expression (***Fig. 5A***). Similar effects of DCMU on NR activity and NO₃⁻ assimilation have been reported in various photoautotrophs [74, 75, 77–79]. However, regulation via PSI, the redox state of the plastoquinone pool, or sugars and their derivatives has also been proposed [74, 80–82]. We therefore suggest that NR synthesis in Symbiodiniaceae depends on nitrate availability, while its stability and activation may depend on photosynthetic electron transport and cellular redox state.

Interestingly, NR gene expression patterns differed among experiments depending on sampling under light or dark conditions (***Fig. 3B,D*** *vs.* ***Fig. 5A***). This likely reflects light-responsive transcription, consistent with reports that dinoflagellate gene expression can vary diurnally even in the absence of strong circadian control [33, 38, 83, 84]. NR transcription may therefore follow a binary “on–off” pattern driven by either light or nitrate exposure, while post-transcriptional mechanisms determine protein abundance.

### NR expression in symbiotic context

Although our culture experiments clarify NR regulation in free-living Symbiodiniaceae, symbionts function within the highly controlled environment of the coral host [85]. We demonstrate that symbionts within three coral species (*S. pistillata*, *T. reniformis*, and *E. lamellosa*) can synthesize NR *in hospite* following nitrate exposure providing the first direct evidence of symbiotic NR induction. Exogenous NO_3_^-^ was rapidly transported through host tissues to the symbionts, where it triggered NR synthesis within 3 h. The absence of NH_4_^+^-mediated repression *in hospite* suggests that intracellular NH_4_^+^ concentrations were insufficient to inhibit enzyme induction under nitrogen-depleted conditions. After 6 h, NR protein abundance declined across all coral species, indicating that NR expression is a transient response to newly available nitrate under nitrogen-depleted conditions, downregulated once nitrate is assimilated to ammonium or nitrogen repletion is restored.

Measurements in FIS further revealed that NR expression in *S. pistillata* symbionts increased in a concentration-dependent manner, consistent with patterns observed in cultured *S. microadriaticum* (***Supplementary Material S10***). In contrast, FIS from *T. reniformis* showed no clear trend, possibly reflecting species-specific nitrogen acquisition strategies [86, 87] that differentially influence the regulation of nitrogen assimilation enzymes. Notably, in FIS, NR protein signals normalized to Cytf and NR-positive controls were substantially higher than those of *in hospite* samples. Although western blotting is semi-quantitative, this consistent pattern suggests that NR expression is lower in symbionts within the host environment. Such repression likely reflects both reduced nitrate availability, due to dependence on host-mediated transport across host and symbiosome membranes, and the presence of alternative nitrogen sources supplied by the host, including NH₄⁺, urea, and amino acids.

Strategies of host-symbiont nitrogen exchange differ among coral species and symbiont lineages. *S. pistillata*, for instance, transfers heterotrophically derived amino acids to its symbionts, whereas *T. reniformis* primarily provides NH_4_^+^ [88]. Corals harbouring *Cladocopium* typically exhibit higher heterotrophic capacity but lower inorganic nitrogen assimilation than those hosting *Symbiodinium* [89], although *Cladocopium* has been shown to assimilate NO_3_^-^ more efficiently than *Durusdinium* [86]. The differences imply that NR expression in symbiotic Symbiodiniaceae is finely tuned to the coral’s nutritional state and to the balance between organic and inorganic nitrogen sources. When organic nitrogen or ammonium is readily available, nitrate assimilation appears to be suppressed, consistent with our observation that NR protein was undetectable in *S. pistillata* symbionts under regular feeding conditions without nitrogen depletion (***Supplementary Material S11***). A key finding of this study is therefore that nitrogen depletion, either in free-living cells or in the coral host, is a prerequisite for detectable NR induction in response to nitrate enrichment.

## Conclusions

Collectively, our results reveal that Symbiodiniaceae possess a tightly regulated, substrate-induced NR system that enables rapid yet transient nitrate assimilation under nitrogen-depleted conditions. NR synthesis occurs within hours of nitrate availability and is actively suppressed by ammonium through post-transcriptional mechanisms. This rapid regulation likely reflects an adaptive strategy to fluctuating nutrient environments, allowing symbionts to exploit transient nitrate pulses without compromising nitrogen homeostasis. Within the coral holobiont, this flexibility provides an auxiliary nitrogen-acquisition pathway, complementing ammonium recycling and heterotrophic nitrogen transfer.

Our findings highlight the importance of post-transcriptional and post-translational control in Symbiodiniaceae nitrogen metabolism and underscore the limitations of transcriptomic data alone for interpreting functional responses. Integrating proteomic and metabolomic approaches [90–94] will be essential to fully elucidate nitrogen regulation within the coral symbiosis and its role in holobiont nutrient balance and resilience.

## Acknowledgements

The authors would like to thank M. Hanikenne and M. Schloesser for their guidance and technical help concerning qPCR work. The authors also thank D. Desgré, M-I. Marcus and E. Elia for their technical help during the experimental setup.

## Author Contributions

CS and SR conceived and designed the experiments with the contributions of RG, CFP, and JCP. CS performed the experiments and analyzed the data. RG and CFP contributed with access to research facility and materials. CS wrote the paper. SR, CFP, RG, and JCP contributed to manuscript revision.

## Data Availability

Data are available via the Zenodo repository: https://doi.org/10.5281/zenodo.18352345

