## Supplementary Information for "Novel Evidence for Nitrogen-Dependent Regulation of Nitrate Reductase in Coral Symbiosis"

### Supporting Information

**S1. Supplementary material and methods.** Identification of Symbiodiniaceae strain identities through ITS2 sequencing.

Symbiodiniaceae strain identities were confirmed by ITS2 sequencing. DNA was extracted from algae culture cell pellets using the DNeasy Plant Mini Kit (QIAGEN) following the manufacturer's instructions and by disrupting the cells in AP1 buffer using acid-washed glass beads (710-1180 µm, Sigma-Aldrich) and a TissueLyser II (QIAGEN) set to 30 Hz for 5 min. DNA concentration and purity were quantified by absorbance measurement on a Synergy Mx Biotek™ Plate Reader using a Take3 Microvolume Plate (Agilent, USA). The Symbiodiniaceae ITS2 gene marker was amplified using forward primer SYM_VAR_5.8S2 5'-GAATTGCAGAACTC-CGTGAACC-3' and reverse primer SYM_VAR_REV 5'-CGGGTTCWCTTGTYTGACTTCATGC-3' [1]. Between 5 and 10 ng of template DNA were used for PCR reaction using the Phusion High-Fidelity PCR Master Mix (Thermo Scientific) with initial denaturing at 98°C for 2 min followed by 35 cycles of 98°C for 10 sec, 56°C for 30 sec and 72°C for 30 sec, and finishing with a final extension at 72°C for 5 min. PCR amplification products were checked on a 2% agarose gel and concentrations were measured on the Synergy Mx before 15 µL of sample at 5 ng/µL were prepared for sequencing using the Mix2Seq kit and Mix2Seq overnight Sanger sequencing services (Eurofins, Germany). Resulting ITS2 sequences were locally blasted on the GeoSymbio database [2] using the NCBI blastn tool [3]. In this way, SSB01 cultures were confirmed as a B1 Symbiodiniaceae strain, corresponding to *Breviolum minutum* (blast score: 481, E-value: $4^{-138}$), and the cultures isolated from *S. pistillata* were confirmed as A1 strain, corresponding to *Symbiodinium microadriaticum* (blast score: 431, E-value: $3^{-123}$).

**Table S2.** Composition of f/2 medium for the different N treatments.

| Component | Stock Solution | Quantity in 1L ASW | Final Molar Concentration |
|---|---|---|---|
| Base for all f/2 media (corresponding to composition of f/2 NF medium) | | | |
| $NaH_2PO_4$ $H_20$ | 5 g/L di$H_2O$ | 1 mL | 36.3 µM |
| Trace metal solution | [see S2 for composition] | 1 mL | – |
| Vitamin solution | [see S3 for composition] | 0.5 mL | – |
| Addition for f/2 $NO_3^-$ 500 µM medium | | | |
| $NaNO_3$ | 75 g/L di$H_2O$ | 0.570 mL | 500 µM |
| Addition for f/2 $NH_4^+$ 500 µM medium | | | |
| $NH_4Cl$ | 26.7 g/L | 1 mL | 500 µM |

**Table S3.** Composition of trace metal stock solution.

| Component | Stock Solution | Quantity in 1L ASW | Final Molar Concentration |
|---|---|---|---|
| $FeCl_3$ $6H_2O$ | – | 3.15 g | $1.17 \times 10^{-5}$ M |
| $Na_2EDTA$ $2H_2O$ | – | 4.36 g | $1.17 \times 10^{-5}$ M |
| $CuSO_4$ $5H_2O$ | 9.8 g/L di$H_2O$ | 1 mL | $3.93 \times 10^{-8}$ M |
| $Na_2MoO_4$ $2H_2O$ | 6.3 g/L di$H_2O$ | 1 mL | $2.60 \times 10^{-8}$ M |
| $ZnSO_4$ $7H_2O$ | 22.0 g/L di$H_2O$ | 1 mL | $7.65 \times 10^{-8}$ M |
| $CoCl_2$ $6H_2O$ | 10.0 g/L di$H_2O$ | 1 mL | $4.20 \times 10^{-8}$ M |
| $MnCl_2$ $4H_2O$ | 180. 0 g/L di$H_2O$ | 1 mL | $9.10 \times 10^{-7}$ M |

**Table S4.** Composition of vitamin solution.

| Component | Stock Solution | Quantity in 1L ASW | Final Molar Concentration |
|---|---|---|---|
| Thiamine HCl (vit. B1) | – | 200 mg | $2.96 \times 10^{-7}$ M |
| Biotin (vit. H) | 0.1 g/L di$H_2O$ | 10 mL | $2.05 \times 10^{-9}$ M |
| Cyanocobalamin (vit. $B_{12}$) | 1.0 g/L di$H_2O$ | 1 mL | $3.69 \times 10^{-10}$ M |

Composition of f/2 detailed in table S1, S2 and S3 are obtained from the literature [4, 5].

**S5.**

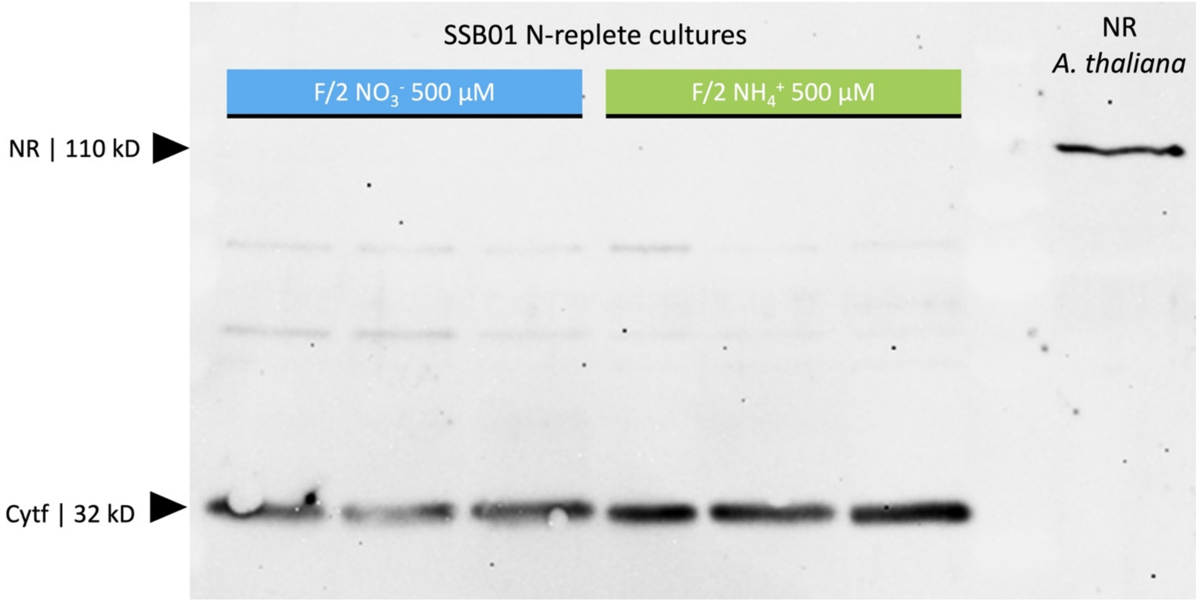


**Figure S5. Western blot analysis showing the failure to detect NR protein from N-replete SSB01 cultures.** Proteins were extracted from N-replete SSB01 grown and maintained on the long-term in f/2 medium with 500 µM of either $NO_3^-$ (blue) or $NH_4^+$ (green). Western blotting for the detection of the NR protein was conducted. The NR protein could not be detected in any of the N-replete SSB01 protein samples as illustrated on the image of one western blot membrane with top row at size ~110 kDa corresponding to NR protein, and a bottom row at size ~32 kDa corresponding to the loading control (protein Cytf). A purified NR from *A. thaliana* served as a positive control.

**S6. Supplementary material and methods.** Analysis and quantification of immunoblot bands using the iBright$^{TM}$ analysis software (Thermo Scientific, USA).

The area of the bands corresponding to nitrate reductase (110 kDa) and cytochrome f (32 kDa) were manually selected, and the volume of each band, corresponding to the sum of each pixel grayscale intensity in the region of the band, was measured and corrected for local background noise (local bg. corr. vol.) by the software. Nitrate reductase protein expression was calculated as the ratio between NR band and Cytf band signal expressed as local background corrected volume.

**S7. Supplementary material and methods.** Selection of primers for housekeeping genes.

Several primer pairs for housekeeping genes (HKG) were selected from the literature (table S7) [6–9] and tested using a PCR profile consisting of an initial denaturation of 2 min at 94°C, followed by 35 cycles of 30 s at 94°C, 30 s at 55°C and 30 s at 72°C, with a final elongation of 7 min at 72°C, using GoTaq® Green Master Mix (Promega, USA). PCR amplification products were visualized on 2% agarose gels using GeneRuler 1kb Plus DNA ladder (Thermo Scientific, USA) for amplicon size estimation. Primers yielding successful amplifications on SSB01 cultures were selected and tested for stability using the geNorm analysis tool in Qbase+ [10, 11]. Based on this, primer pairs Cyc_Maru [9] and Act1_May [8] for genes cyclophilin and actin1, respectively, were selected as HKG for all experiments investigating NR gene expression levels.

**Table S7.** List of tested HKGs with references from the literature

| Gene | Primer pair name | Target type | Forward primer (5'-3') | Reverse primer (5'-3') | Amplicon size | Amplification test | qPCR & geNorm test | Reference |
|---|---|---|---|---|---|---|---|---|
| Calmodulin | Cal_Kru | B1 | TGATGGCGCGCAAGATGAAGG | TGCCATCGCGATCGAAAACCTTG | 78 bp | Passed | Failed melt curve | (Krueger et al. 2015) |
| Cytochrome Oxidase Subunit 1 | Cox_Kru | B1 | TCTGTCTTCCTCTCACATCTCT | CCACTGCACCATTTCCAAGA | 82 bp | Passed | Rejected by geNorm test | (Krueger et al. 2015) |
| S4 Ribosomal Protein | RpS4_Ros | C3 | CCGCACAAACTGCGTGAGT | CGCTGCATGACGATCATCTT | 101 bp | Passed | Failed melt curve | (Rosic et al. 2011) |
| Actin | Act1_May | B1 | CACCACCATGTTCACAGGAA | AGCCACCACCTTGATCTTCA | 87 bp | Passed | Passed | (Mayfield et al. 2014) |
| Cyclophilin | Cyc_Ros | C3 | ATGTGCCAGGGTGGAGACTT | CCTGTGTGCTTCAGGGTGAA | 101 bp | Passed | Failed melt curve | (Rosic et al. 2011) |
| Cyclophilin | Cyc_Maru | B1 | TGCTTGAGGGTAAAGTTCTC | CAACCCCTTCATTTCAAAGG | N.A. | Passed | Passed | (Maruyama et al. 2022) |

**S8. Supplementary material and methods.** Quantification of NR gene expression from Ct values.

Ct values (threshold cycle with fluorescence threshold set to 0.2) of technical triplicates were averaged for each sample, and the Ct values of the two HKGs Cyc_Maru and Act1_May were averaged and subtracted from the mean Ct value of the sample to obtain ΔCt ($\Delta Ct = Ct_{sample} - (Ct_{Cyc\_Maru} + Ct_{Act1\_May})/2$). The ΔCt value was then subtracted from the mean ΔCt value of the samples at time 0 h to express relative expression ΔΔCt. Fold gene expression was calculated as $2^{-\Delta\Delta Ct}$ for each sample (biological replicate) in each experimental condition.

**S9. Supplementary material and methods.** *In hospite* experimental design and set up.

All coral colonies were maintained in flow-through seawater aquaria at the Centre Scientifique de Monaco (Monaco). One mother colony of each species was used to produce 36 nubbins which were evenly distributed and allowed to recover in two independent aquaria (28 L) supplied continuously with natural oligotrophic seawater (0.5 µM DIN and 0.2 µM DIP) at a flow rate of 20 L $h^{-1}$, pumped at a depth of 50 m. During the four weeks of recovery, nubbins were fed twice a week with *Artemia* nauplii, maintained at 25°C ± 0.5°C, with constant agitation provided by water pumps, and under 150 µmol photons $m^{-2}$ $s^{-1}$ in a 12h:12h day:night cycle. Feeding of the coral nubbins was stopped two weeks prior to the experiments to avoid interaction with the nitrogen enrichments.

All treatments were carried out in duplicate 28 L tanks under similar temperature (25°C ± 0.5 °C), flow, and light (150 µmol photon $m^{-2}$ $s^{-1}$) conditions as described above. For both experiments, coral nubbins were first incubated in nitrogen-depleted water for 24 h (undetectable levels of dissolved inorganic nitrogen, DIN). All experimental tanks were kept as a closed system and nitrate enrichment was achieved by mixing a $NaNO_3$ concentrated stock solution to the tank seawater just before coral nubbins were transferred to the experimental tanks.

For experiment B, symbionts were freshly isolated (FIS) from the coral host tissues using waterpick blasting with 0.2 µm filtered seawater (FSW). Coral tissue slurry was then homogenized using a syringe equipped with a 24G needle to dissociate symbiont cells from the coral tissue, followed by three centrifugation and wash steps to pellet symbiont cells and remove the supernatant. FIS were incubated in 200 mL culture flasks with ventilated cap, at a temperature of 25°C ± 0.5 °C with gentle agitation using a magnetic stirrer and under 150 µmol photons $m^{-2}$ $s^{-1}$ constant light.

**S10.**

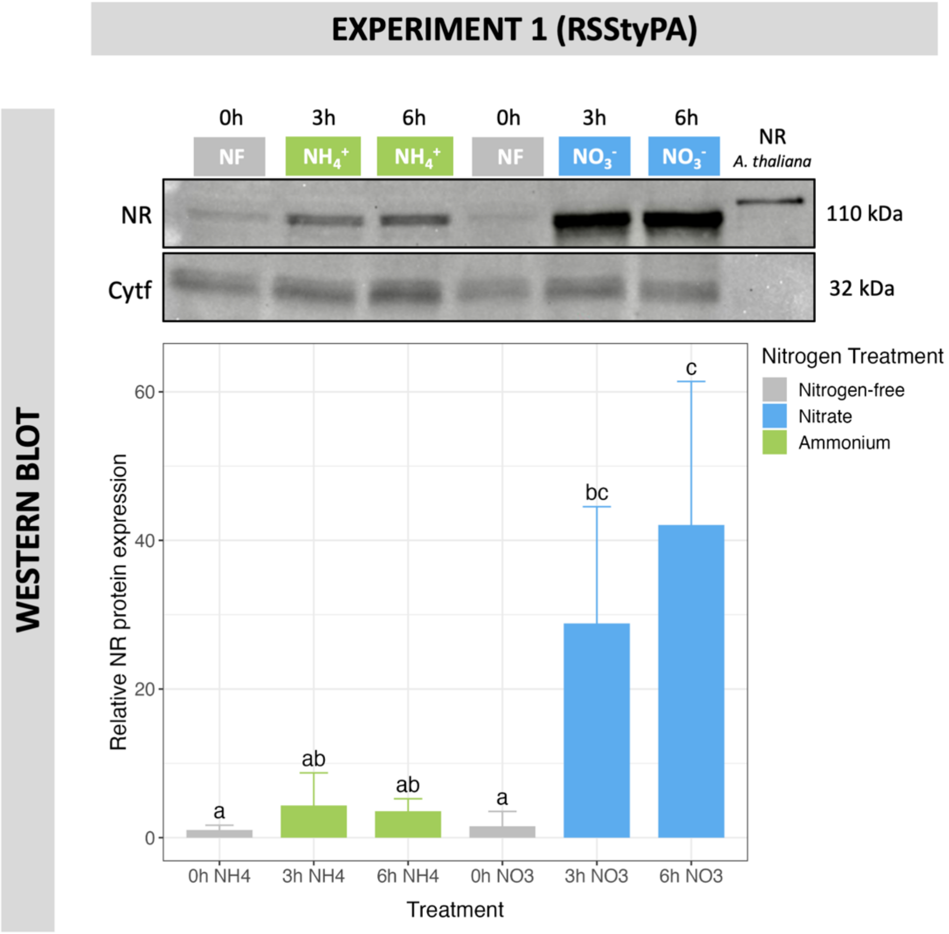


**Figure S10. Kinetics of NR expression in cultured *S. microadriaticum*: WB results from Experiment 1.** Incubation of N-depleted algae in f/2 $NO_3^-$ 500 µM or f/2 $NH_4^+$ 500 µM for 3 and 6 hours. Only one WB membrane (out of 3 replicates) is displayed on this figure as a representation, with a top row at size ~110 kDa corresponding to NR protein, and a bottom row at size ~32 kDa corresponding to the loading control (protein Cytf). Mean relative signals of WB bands from all replicates (n=3) are represented on the bar plot. Statistically significant differences are represented by different letters (a, b and c). Error bars correspond to the 95% confidence interval (CI).

**S11**

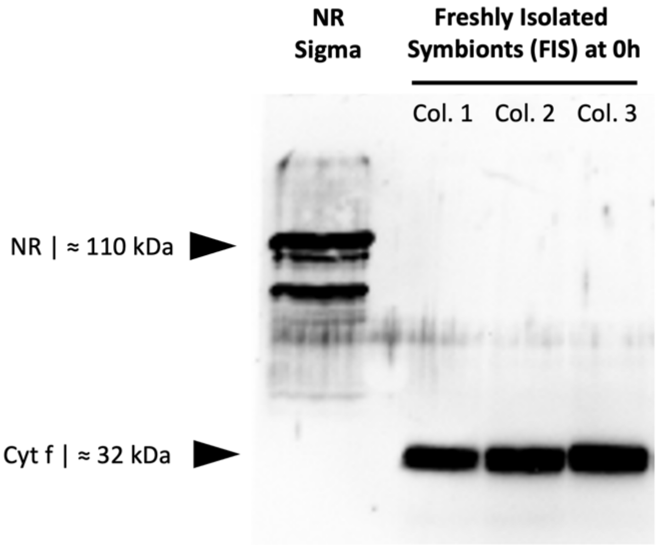


**Figure S11. Western blot analysis showing failure to detect NR protein in FIS from N-replete corals.** After incubation of N-replete *S. pistillata* corals in seawater enriched with $NO_3^-$, symbionts were isolated (FIS) and their proteins extracted for western blotting for the detection of the NR protein. The NR protein could not be detected in any of the FIS samples as illustrated on the image of one western blot membrane with top row at size ~110 kDa corresponding to NR protein, and a bottom row at size ~32 kDa corresponding to the loading control (protein Cytf). NR Sigma is a purified NR from *A. thaliana* serving as a positive control.
